# GC-MS Profiling of Compounds produced by endophytic fungi ex-situ and from their host plants, *Azadirachta indica* and *Melia azedarach* collected in Kenya, Africa

**DOI:** 10.64898/2026.04.16.719096

**Authors:** Rita Dill, Truphosa Amakhobe, Grace Oballa, George Ojenge, Favour Adibe, Jiangnan Peng, Sheila Okoth, Anne Osano

## Abstract

Endophytic fungi residing within medicinal plants are emerging as prolific sources of structurally diverse bioactive secondary metabolites with applications in drug discovery. *Azadirachta indica* (Neem) and *Melia azedarach* (Melia), members of the *Meliaceae* family, are renowned for their rich phytochemical composition; however, the contribution of their endophytic fungi communities to this chemical diversity remains largely unexplored. Herein, endophytic fungi were isolated from leaves and bark of Neem and Melia collected in Kenya and cultured under distinct physical conditions, solid (plates) and liquid (broth) media to assess how culture environment influences compound production. Compounds were extracted and analyzed using gas chromatography-mass spectrometry (GCMS) to profile the chemical diversity associated with each endophytic fungi, physical culturing state and host plant. GCMS analysis revealed that while the host plant identity influences the presence of specific compounds, the dominant determinant of chemical diversity was intrinsic biosynthetic capacity of the endophytic fungi themselves. Several compounds were unique to endophytic fungi cultures, highlighting their role as independent sources of bioactive compounds. Culture conditions moderately influence metabolite profiles, demonstrating the importance of optimizing growth environments in experimental design and natural product bioprospecting. From the Neem samples, we found 53 compounds uniquely present in the broth samples (consisting of Neem powder and endophytic fungi), 22 found exclusively with the endophytic fungi from the Neem, and 31 compounds shared between the broth and the endophytic fungi samples. In Melia samples, 109 compounds were uniquely present in broth samples from Melia plant (consisting of Melia powder and endophytic fungi), 22 compounds were found exclusively with the endophytic fungi from the Melia, and 55 were shared between the broth and the endophytic fungi samples. Our comparative analysis assessed the Neem and Melia endophytic fungi exclusive samples and reported 12 shared compounds. 10 compounds were unique to Neem and 10 unique to Melia; however, their identities varied between the two categories. While GCMS enabled the identification of volatile and semi-volatile metabolites, future studies employing complementary metabolomic approaches, such as liquid chromatography-mass spectrometry (LCMS), ultra-high-performance liquid chromatography MS/MS (UHPLC MS/MS), or nuclear magnetic resonance (NMR) spectroscopy, would expand coverage to non-volatile, polar, and high molecular weight compounds, providing a more comprehensive understanding of endophyte-derived chemical diversity. These findings provide insights into the interplay between medicinal plants and their endophytes and establish a foundation for leveraging endophytic fungi from Neem and Melia as scalable sources of structurally complex natural products for pharmaceutical and biotechnological applications while minimizing ecological impact.

## Introduction

Endophytic fungi are ubiquitous microorganisms that inhabit plant tissues without causing harm and are increasingly recognized as prolific sources of bioactive secondary metabolites; these metabolites are often broadly termed and will be referred to as such within our writings [1,4,7,8]. These compounds often exhibit antibacterial, antifungal, antiviral, and anti-inflammatory activities, making endophytic fungi promising targets for natural product discovery and pharmaceutical development [2,3,5,9]. Therefore, exploiting endophytic fungi offers a sustainable alternative to harvesting whole plants, enabling rapid access to bioactive compounds while minimizing ecological impact and supporting biodiversity conservation [16,17,10].

*Azadirachta indica* (neem) and *Melia azedarach* (melia), members of the Meliaceae family, are widely used in traditional medicine across Asia, Africa, and other tropical regions [18,19,11]. They are rich in structurally diverse compounds, including limonoids, terpenoids, and phenolic compounds, which serve as scaffolds for drug development [20,21]. Limonoids, highly oxygenated triterpenoids such as azadirachtin, nimbin, salannin (in Neem), and meliatoxins or melianones (in Melia), exhibit potent insecticidal, antiviral, and anticancer activities [22,23,13]. Terpenoids, including mono-, sesqui-, and diterpenes, contribute to aromatic and medicinal properties, while phenolics and flavonoids, such as quercetin, rutin, and catechins, provide antioxidant, antibacterial, and anti-inflammatory effects [20,12]. Collectively, these metabolites underpin the medicinal versatility of neem and melia and make them attractive targets for natural product research [20-23].

While plant-derived compounds are valuable, their extraction often requires large-scale harvesting, which may threaten local biodiversity [24-26]. Endophytic fungi associated with these plants present a complementary approach, as they can be isolated, cultured, and manipulated *in vitro*, providing faster and more sustainable access to complex bioactive compounds [7,9]. Furthermore, endophytic fungi may produce metabolites that are like, or distinct from, those of their host plants, expanding the chemical diversity accessible for drug discovery [8,10]. Accurate characterization of metabolite origin is therefore critical to distinguish plant-derived from endophyte-derived compounds and to guide bioprospecting strategies that minimize ecological impact [27-29].

Within this work we have collected Neem and Melia leaf tissues from Kenya, Africa. We cultured the endophytic fungi from the leaves of each medicinal plant in solid (plates) and liquid (broth) physical states to determine if the different physical culturing states impact the presence or absence of compounds produced by the endophytic fungi. Finally, we compared if the host plants affected the compounds produced by the endophytic fungi. The resulting compounds extracts were tested using gas chromatography-mass spectrometry (GC-MS) to note the extent by which endophytic fungi, the physical culturing states, and the host plant used would alter the presence or absence of compounds or their overall profiles. We hypothesized that some compounds would be unique to endophytic fungi, highlighting their biosynthetic potential, and that while culture conditions may modulate metabolite profiles, the intrinsic diversity of the endophytic community would remain the primary driver of chemical variation. By demonstrating how endophytic fungi can provide rapid, sustainable access to structurally complex metabolites, this work supports biodiversity-conscious approaches to natural product discovery and lays the foundation for exploiting endophytic fungi as scalable sources of novel compounds for pharmaceutical and biotechnological applications.

## 2. Materials and Methods

### 2.1 Sample collection

Leaves and bark of *Azadirachta indica* (Neem) and *Melia azedarach* (Melia) were collected from healthy, mature trees in Machakos County, Kenya during the month of June in the year of 2026. From the Neem and Melia trees leaves were collected. All samples were handled with sterile gloves and placed in sterile polyethylene bags to prevent cross-contamination. Samples were transported to the laboratory on ice and processed within 24 hours. Voucher specimens were deposited at Nairobi University Herbarium (NAI).

### 2.2 Isolating and Culturing Endophytes on Plates

The Neem and Melia whole plant tissue was rinsed with tap water for approximately 30 s to remove surface debris. Using a clean surgical blade, leaves were cut into approximately 1 cm long pieces on the workbench. The pieces were placed into empty Petri dishes. The Petri dishes were filled with enough 95% ethanol to lightly cover plant pieces and then agitated for 10 s before being drained. Next, the dish was filled with 10% Clorox and agitated for 2 min, then drained. The dish was then filled with 70% Ethanol and agitated for another 2 min, then drained. The dish was then opened slightly in the hood and left to dry. Flame-sterilized and cooled forceps were used to transfer the surface-sterilized plant pieces onto Malt Extract Agar or Potato Dextrose Agar. The plant pieces were spaced to avoid overgrowth, and the plates were wrapped with parafilm. The plates were clearly labelled and incubated at room temperature for 7 days. After fungal growth emerged, the colonies were subcultured onto freshly prepared media to obtain pure cultures.

### 2.3 Extracting Endophyte Compounds from Solid Media Culture

After sub-culturing, a total of 24 pure fungal isolates were obtained. Each isolate was sub-cultured onto freshly prepared media to obtain sufficient biomass for compounds extraction. The cultures were then incubated for 7 days at room temperature. Following incubation, fungal mycelia and associated agar were aseptically scraped in a laminar flow hood using sterile scalpel blades and then placed in a 50 mL Falcon tube. 25 mL of ethyl acetate (EtOAc) was then added to each Falcon tube. A clean, autoclaved wooden tongue depressor was used to push any endophyte samples off the walls of the tube and into (EtOAc) and to smash the sample, disrupting the mycelium, which improves extraction. The mixture was vortexed to initiate compounds dissolution. As an alternative to sonicating, the tubes were then placed lying flat in an orbital shaker and agitated for 1 hr at 240 rpm to enhance extraction. Subsequently, the mixtures were centrifuged at 3000 rpm for 15 min to separate the organic and aqueous phases. The supernatant was carefully collected, and the extraction process was repeated once on the remaining pellet to maximize compounds recovery. The combined supernatant was concentrated using a rotary evaporator under reduced pressure at 40°C. The extracts were placed in sterile vials at 4°C pending further analysis. These extracts were later subjected to GC-MS analysis for compounds profiling

### 2.4 Extracting Endophyte Compounds from Liquid Broth

Potato Dextrose Broth (PDB) was prepared and autoclaved, then cooled to approximately 45°C. Neem or Melia powder corresponding to the source of each fungal isolates, was added to the medium. For each culture, 200 mL of supplemented PDB was dispensed into a sterile conical flask, and a single fungal isolate was inoculated into each flask under aseptic conditions. Control flask contained PDB with Neem and Melia powder but no fungal inoculum. The flasks were incubated at room temperature (∼25 °C) for 7 days on an orbital shaker set at 240 rpm. After incubation, the cultures were gently macerated using sterile, autoclaved wooden tongue depressors to disrupt mycelial mats, and the broth was filtered through Whatman paper to remove fungal biomass. The filtrate (15-25 mL per sample) was collected into sterile 50 mL Falcon tubes. An equal volume of (EtOAc) was added to each filtrate (1:1 ratio), and the mixtures were vortexed for 10 s to initiate metabolite dissolution. The tubes were then placed lying horizontally in an orbital shaker and agitated at 240 rpm for 1 hr. The mixtures were then centrifuged at 3000 rpm for 15 min to separate the organic and aqueous phases. The supernatant was collected, and the extraction procedure was repeated once on the residual aqueous phase to maximize metabolite recovery. The combined organic extract were concentrated under reduced pressure using a rotary evaporator at 40°C impact [6]. Concentrated extracts were transferred into sterile vials and stored at 4°C until further analysis.

### 2.5 GC-MS: Gas Chromatography Mass Spectrometry

Samples in EtOAc were allowed to evaporate to dryness under a fume hood and subsequently reconstituted in 500 µL of GC-grade dichloromethane (DCM) (Sigma-Aldrich, St. Louis, MO, USA). The solutions were vortexed for 10 s, sonicated for 10 min, and centrifuged at 14,000 rpm for 5 min. The resulting supernatant was dried over anhydrous Na2SO4, and an aliquot (1.0 µL) was analyzed by GC-MS using a 7890A gas chromatograph (Agilent Technologies, Inc., Santa Clara, CA, USA) coupled to a 5975C mass selective detector (Agilent Technologies, Inc., Santa Clara, CA, USA). The GC system was equipped with a low-bleed (5%-phenyl)-methylpolysiloxane HP-5MS capillary column (30 m × 0.25 mm i.d., 0.25 µm film thickness; J&W, Folsom, CA, USA). Helium was used as the carrier gas at a constant flow rate of 1.25 mL/min. The injector temperature was maintained at 270 °C, while the transfer line temperature was set at 280 °C. The oven temperature program was as follows: initial temperature of 35 °C held for 5 min, increased at 10 °C/min to 280 °C, and held for 20.4 min. The mass selective detector was operated under electron impact (EI) ionization at 70 eV, with the quadrupole and ion source temperatures maintained at 180 °C and 230 °C, respectively. Mass spectra were acquired in full-scan mode over an m/z range of 40-550, with a solvent delay of 3.3 min. Blank runs of the reconstituting solvents (DCM and EtOAc) and instrument blanks were analyzed under identical conditions, and their corresponding peaks were excluded from the analysis. Compound identification was based on comparison of retention times and mass fragmentation patterns with those of authentic reference standards where available, as well as reference spectra from the National Institute of Standards and Technology (NIST) mass spectral libraries (versions 05, 08, and 11).

### 2.6 Principal Component Analysis (PCA) and presence versus absence bar charts

PCA was executed using presence-absence GC-MS data based upon identified peak area. Compound data was converted to binary format where 1 indicated the presence of a compound and 0 indicated an absence thereof. This in turn yielded our samples as observations and our compounds as variables. PCA was performed using the PCA implementation from the scikit-learn library (v.1.x) within Python. We did not apply scaling or transformation prior to analysis. Samples were plotted in 2-D ordination based on their PCA scores. The plotting was annotated and colored according to the experimental conditions.

### 4. Results

We analyzed our GC-MS findings further using PCA and presence-absence bar charts. This was done in efforts to monitor the influence of culturing conditions, and endophytes from which compounds were extracted on GC-MS profiles. Collectively, our PCA results (Figures 1-4) suggest that culturing in broth or liquid can have a minor influence on the compounds composition. Our GC-MS data (Tables 1-3) simultaneously informs that endophytes can contribute to the unique compounds seen in medicinal plants beyond those made by the plants themselves. The continuous clustering of endophytic fungi alone from plate and broth conditions resonates that the intrinsic endophytic fungi is the primary driver of separation rather than the physical culturing states or the Neem versus Melia host plants. Additionally, the many unique to endophytic fungi compounds, noted in our tables (Tables 1-3), supports the rationale that endophytes are independent contributors to the overall metabolomic profiles of Neem and Melia.

**Figure 1.**
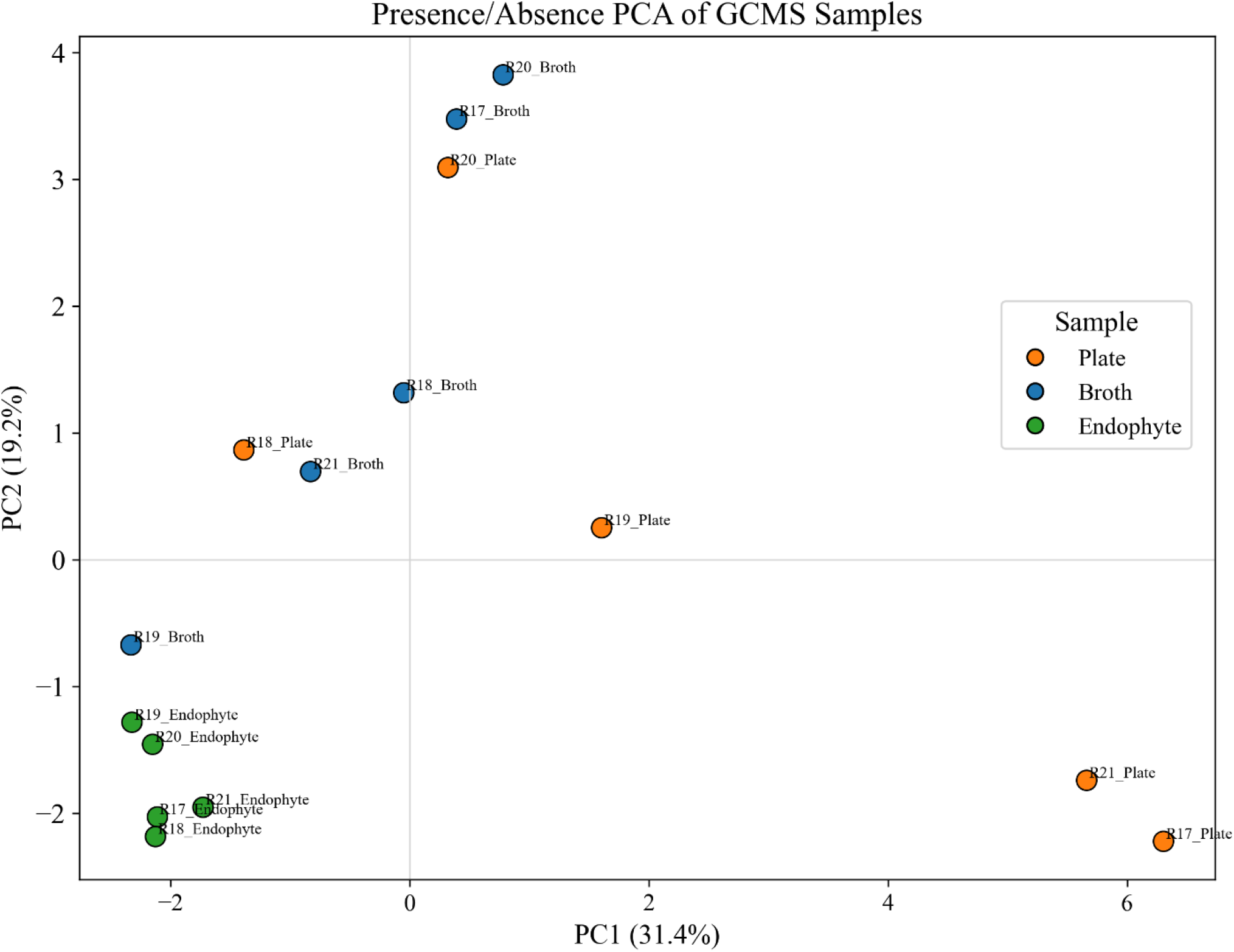
PCA of Neem GC-MS profiles from the broth (blue colored and consisting of endophyte and Neem tissue), plate (orange colored and consisting of endophyte and Neem tissue) and its corresponding endophyte (green colored and consisting of only endophyte) samples. Where “R” denotes an individual endophyte isolate. We find that the endophyte samples cluster independently while the plate and broth samples exhibit little separation.

**Figure 2.**
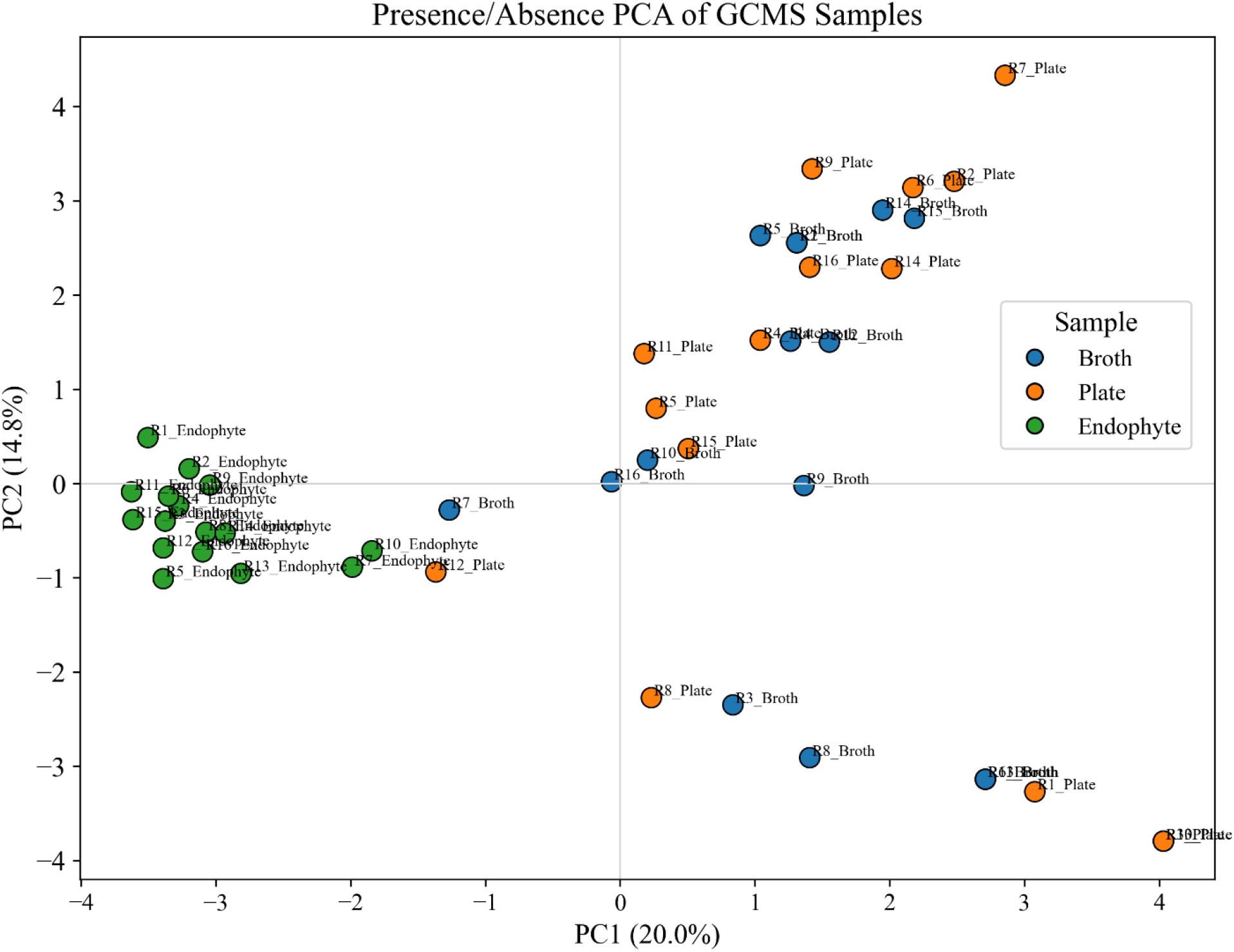
PCA of Melia GC-MS profiles from the broth (blue colored and consisting of endophyte and Melia tissue), plate (orange colored and consisting of endophyte and Melia tissue) and its corresponding endophyte (green colored and consisting of only endophyte) samples. Where “R” denotes an individual endophyte isolate. The PCA shows endophyte samples clustering singly and in contrast the plate and broth samples do not display separation. From here we see that physical growth state minimally influences compounds detected, and the endophyte diversity predominantly drives separation of samples.

**Table 1.** A comparison of the endophytes found on Melia host plants contrasted with the endophytes found on the Neem host plants. This showcases that host specificity may affect the endophyte’s compounds profiles seen.

| **Shared (Neem & Melia endophytes)** | **Melia endophyte** | **Neem endophyte** |
| --- | --- | --- |
| (3E)-Octadecene | (2E)-Pentenal | 2,6,10,14-tetramethyl hexadecane |
| 2,6,10,15-tetramethyl heptadecane | 2,6,11,15-tetramethyl hexadecane | 2,6,10-trimethyl pentadecane |
| 2-Nonanol | 2-Methyl-2-docosene | 2,6,11-trimethyl dodecane |
| 2-methyl-heptadecane | 2-methoxy-3-methyl phenol | 3-methylbutyl butanoate |
| 3-hydroxypropyl octadecanoate | 2-methyl-, 3-methylbutyl propanoate | 6-Tridecanone |
| 5-Decanone | 3,5-dimethoxy phenol | Dodecane |
| 6-Dodecanone | 9,19-Cyclolanost-25-en-3-ol, 24-methyl-, (3.beta.,24S) | Eicosane |
| Ethyl Oleate | Heptacosane | Ethyl propanoate |
| Geranyl isovalerate | Methyl -9Z-hexadecenoate | Isopropyl myristate |
| Hentriacontane | Octadec-9-enoic acid | Oleic Acid |
| Methyl pentadecanoate |  |  |
| Octacosanol |  |  |

## Discussion and Conclusion of Results

The work presented herein has investigated the extent by which culturing conditions, plant tissue, and host plant can drive variation in the profiles of compounds extracts from endophytes found on Neem and Melia. While Neem and Melia have been heavily researched, to the best of our knowledge, our work is novel in that we have systematically compared the GC-MS profiles of the endophytic fungi found on Neem and Melia and modulated the physical state of culturing. Previous work has independently assessed endophytes from Neem and Melia through the lens of diversity or bioactivity. Our PCA analysis of the GC-MS results showcases that compound composition remains moderately conserved across the physical culturing states monitored as noted by their overlap (Figures 3-4). Each sample analyzed, and denoted with “R” in each figure represented a distinct endophyte isolate. Therefore, we believe the intrinsic differences in the compounds between the isolates would be the major driver of variation observed in the PCA. This conclusion is evident through our PCA’s which possess control presence (Figures 1-4). We additionally saw that host plants did not render distinct separation between the broth profiles when comparing Neem and Melia grown with host plant tissue (Figures 5-6). This aligns well with anecdotal reporting of the two being used interchangeably in traditional medicine. Our presence-absence tables (Tables 1-2) were critical in efforts to deepen our understanding of the distinct role endophytes play in compounds composition, and how they may vary across different host plants (Table 3). Due to the microscopic nature of endophytes, it is common for endophytes and their self-produced compounds to be consumed during ayurvedic practices. However, in the pursuit of providing knowledge for drug development by others in the community, it is imperative to identify the true producer of the small compounds. Our presence-absence tables served the vital purpose of such identification. From the Neem samples, we found 53 compounds uniquely present in the Neem tissue with endophyte, 22 found exclusively with the Neem-relevant endophyte, and 31 compounds shared between the two conditions. In Melia samples, 109 compounds were uniquely present in Melia tissue with endophyte, 22 compounds were found exclusively with the Melia-relevant endophyte, and 55 were shared between the two conditions. Our comparative analysis assessed the Neem and Melia endophyte exclusive samples and reported 12 shared compounds. 10 compounds were unique to Neem and 10 unique to Melia; however, their identities varied between the two categories. This further confirms the rationale that endophytes may operate as independent sources of compounds diversity. Furthermore, endophytes may be potential reservoirs of compounds whose structures could have use in efforts to develop new pharmaceutical products.

**Figure 3.**
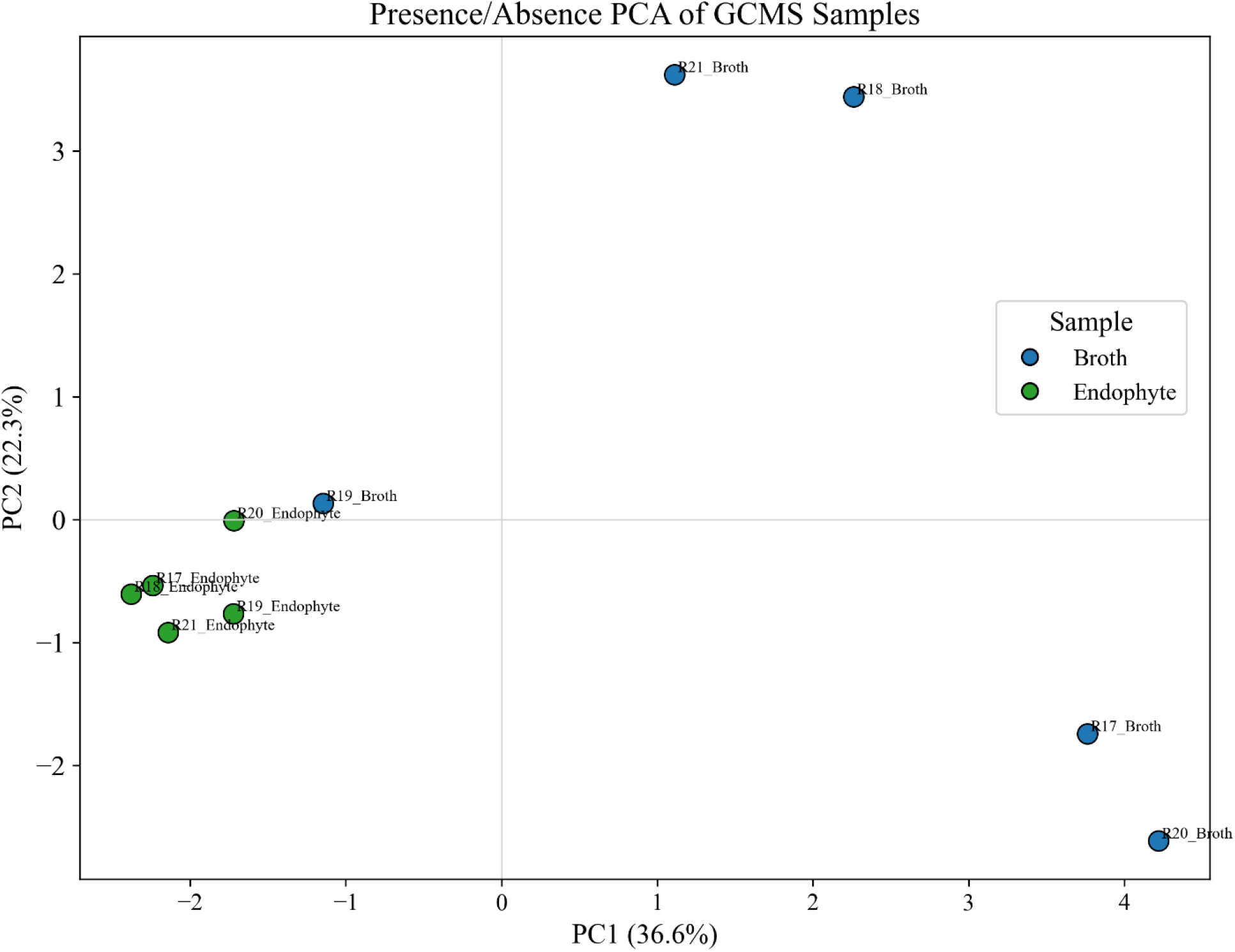
PCA of Neem GC-MS profiles from the broth (blue colored and consisting of endophyte and Neem tissue) and its corresponding endophyte (green colored and consisting of only endophyte isolate) samples. Where “R” denotes an individual endophyte isolate. The broth and control samples separately primarily along the PCA and thus they signal that the endophytes and host tissue compounds profiles have significant content differences.

**Figure 4.**
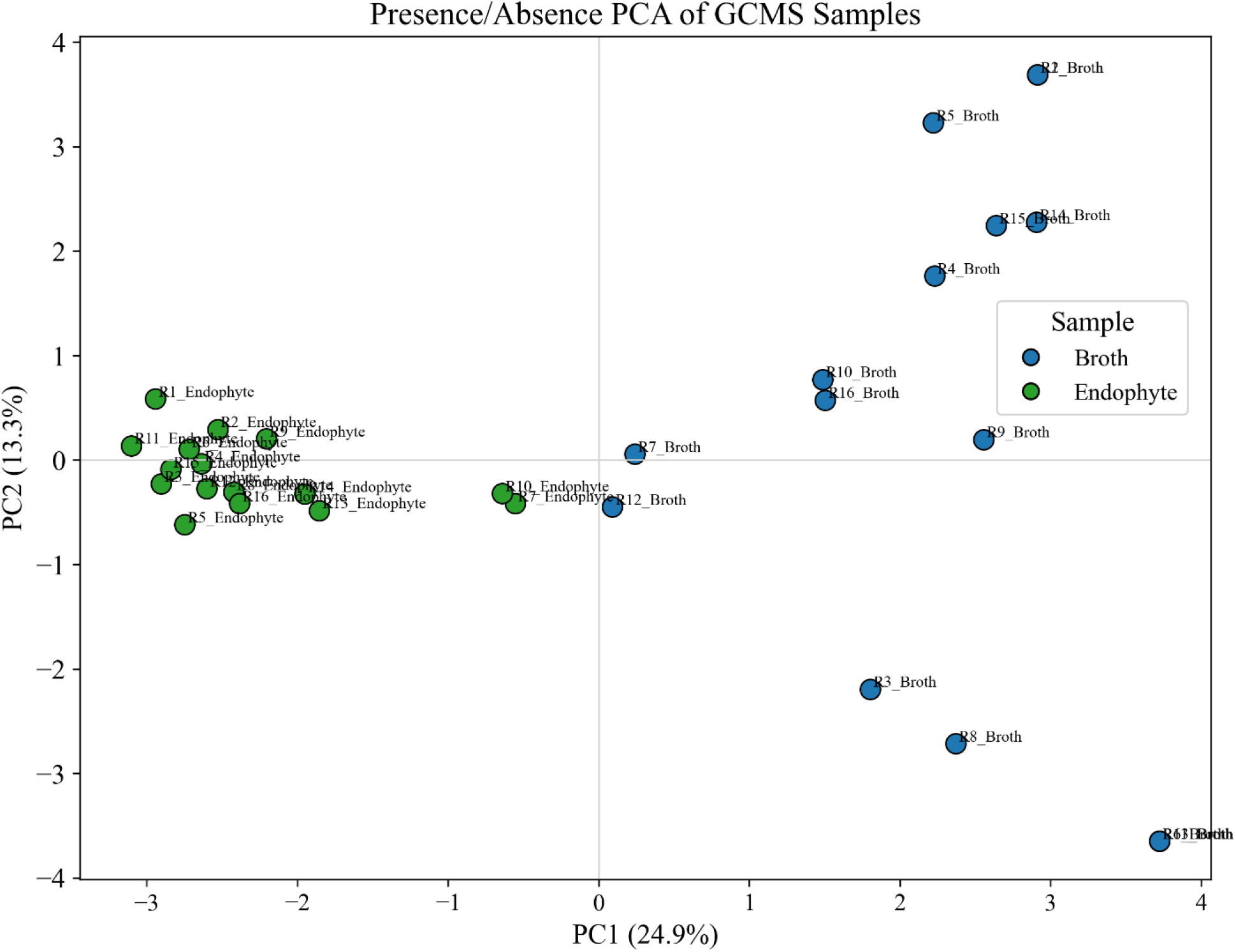
PCA of the endophytic fungi synthesized compounds from the Melia plant. GC-MS profiles from the broth (blue colored and consisting of endophyte and Melia tissue) and its corresponding endophyte (green colored and consisting of only endophyte) samples. Where “R” denotes an individual endophyte isolate. The broth and endophyte samples segregate into groupings and echo the findings seen in the Neem PCA: endophyte and host tissue compounds profiles have significant content differences.

**Figure 5.**
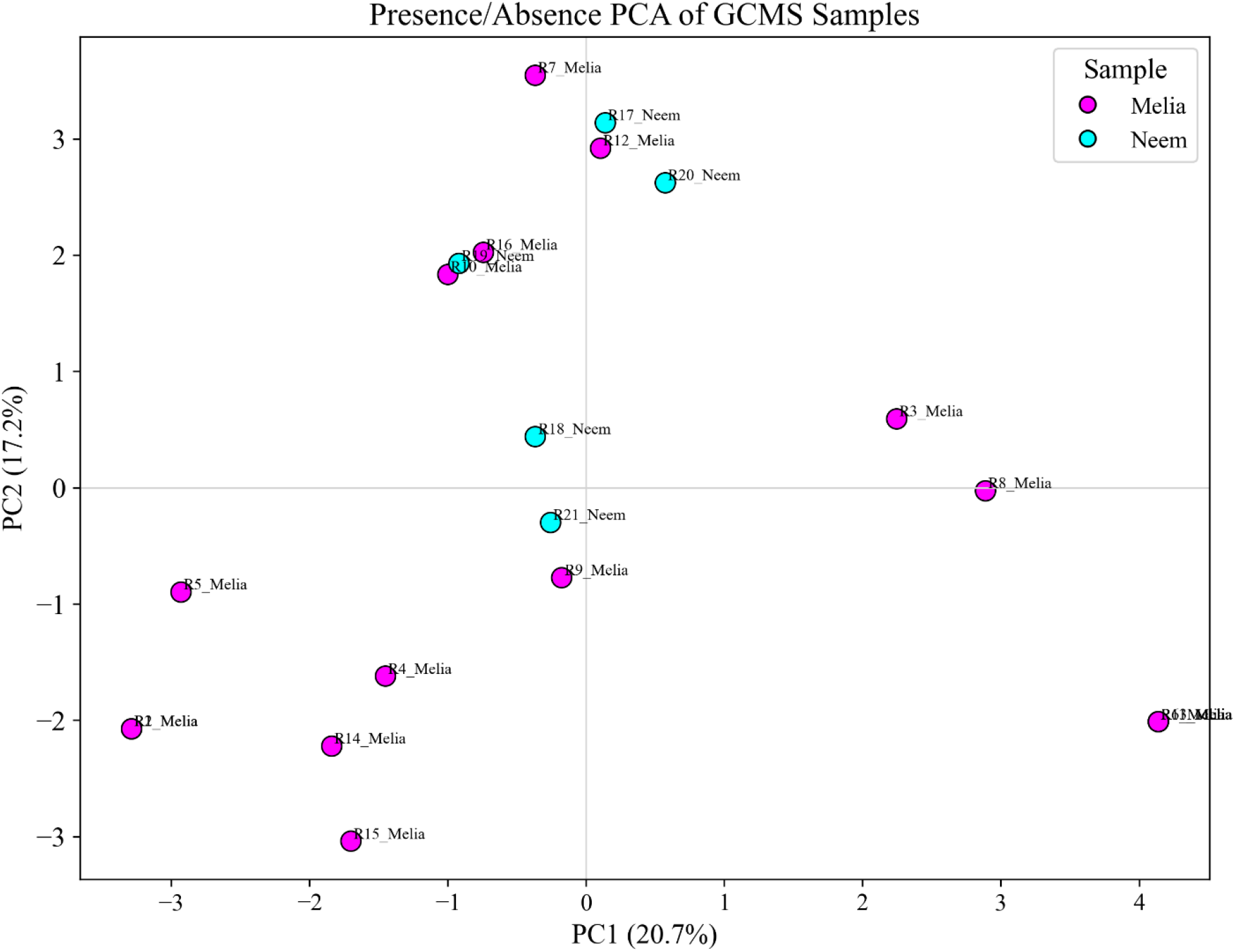
PCA of Melia and Neem GC-MS profiles from the broth (magenta colored is consisting of Melia endophyte and Melia tissue. Cyan colored is consisting of Neem endophyte and Neem tissue) . Where “R” denotes an individual endophyte isolate. The PCA shows that in the case of samples cultured in broth states, the two host plants do not have clear separation.

**Figure 6.**
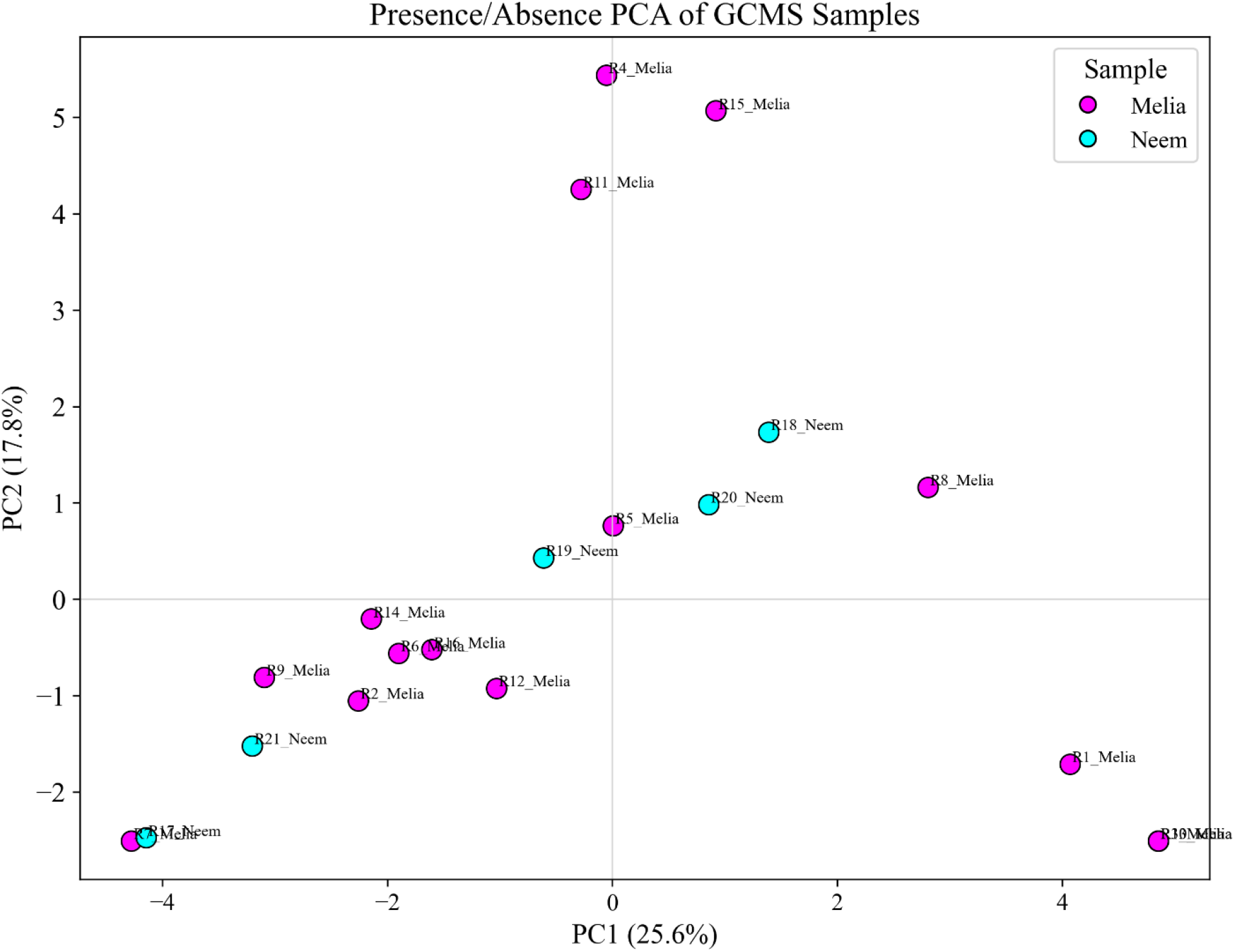
PCA of Melia and Neem GC-MS profiles from the plate (magenta colored is consisting of Melia endophyte and Melia tissue. Cyan colored is consisting of Neem endophyte and Neem tissue) . Where “R” denotes an individual endophyte isolate. The PCA shows little separation between the Melia and Neem profiles. This suggests that the physical state of plate has a small effect on the compounds profiles between the plate-specific samples.

**Table 2.** Compounds detected in Neem and its corresponding control samples. From here it is evident that endophytes create their own compounds, and that their compounds are different from those made by the host plant.

| **Neem + endophyte** | **Endophyte** | **Shared** |
| --- | --- | --- |
| (2E)-Pentenal | (3E)-Octadecene | 1-Docosene |
| (E)-Caryophyllene | 2,6,10,14-tetramethyl hexadecane | 1-Hexadecene |
| 1,2,3,4-Butanetetrol, [S-(R*,R*)]- | 2,6,10,15-tetramethyl heptadecane | 1-Octadecene |
| 1,3-bis(1,1-dimethylethyl)-benzene | 2,6,10-trimethyl pentadecane | 11-Methyldodecanol |
| 1-Decene | 2,6,11-trimethyl dodecane | 2,4-bis(1,1-dimethylethyl) phenol |
| 1-Heptadecene | 2-Nonanol | 2,6,10,14-tetramethyl pentadecane |
| 1-Hexacosene | 2-methyl-heptadecane | 7,9-Di-tert-butyl-1-oxaspiro(4,5)deca-6,9-diene-2,8-dione |
| 13Z-Docosenamide | 3-hydroxypropyl octadecanoate | C20-ol |
| 2,3-Butanediol | 3-methylbutyl butanoate | Docosane |
| 2,3-Butanediol, [R-(R*,R*)]- | 5-Decanone | Eicosane |
| 2,6,10-trimethyl tetradecane | 6-Dodecanone | Ethyl (9E)-octadecenoate |
| 2-hydroxy-1-(hydroxymethyl)ethyl esterh exadecanoic acid | 6-Tridecanone | Ethyl hexadecanoate |
| 2-methyl tridecane | Dodecane | Heneicosane |
| 2-methyloctacosane | Eicosane | Heptacosane |
| 3,7-dimethyl-decane | Ethyl Oleate | Heptadecane |
| 3-Penten-2-ol | Ethyl propanoate | Hexacosane |
| 5-Tetradecanol | Geranyl isovalerate | Hexadecane (C16) |
| 8,11,14-Eicosatrienoic acid, (Z,Z,Z)- | Hentriacontane | Isopropyl tetradecanoate |
| 8,11,14-Eicosatrienoic acid, methyl ester | Isopropyl myristate | Nonadecane |
| 9,12-Octadecadienoic acid (Z,Z)- | Methyl pentadecanoate | Octacosane |
| 9-Octadecenamide, (Z)- | Octacosanol | Octadecane (C18) |
| 9-Octadecenoic acid, (E)- | Oleic Acid | Octadecanoic acid |
| Benzyl alcohol |  | Pentacosane |
| Butanedioic acid |  | Pentadecane |
| Butanoic acid |  | Squalene |
| C14-ol Tetradekanol |  | Tetracosane |
| Dodecane |  | Triacontane |
| Eicosanoic acid |  | Tricosane |
| Ethyl 9-hexadecenoate |  | n-Hexadecanoic acid |
| Ethyl octadecanoate |  | n-Pentadecanol |
| Heptadecanoic acid |  | n-Tetracosanol-1 |
| Hexanoic acid |  |  |
| L-Lactic acid |  |  |
| Limonene |  |  |
| Methyl 9Z-Octadecenoate |  |  |
| Methyl dodecanoate |  |  |
| Methyl octadecanoate |  |  |
| Methyl tetradecanoate |  |  |
| Nonanal |  |  |
| Pentadecanal |  |  |
| Pentadecanoic acid |  |  |
| Phenol |  |  |
| Styrene |  |  |
| Tetracosane |  |  |
| Tetradecane |  |  |
| Tetradecanoic acid |  |  |
| Tridecane |  |  |
| Undecanoic acid |  |  |
| Z,E-2,13-Octadecadien-1-ol |  |  |
| cis-10-Heptadecenoic acid |  |  |
| n-Decanoic acid |  |  |
| n-Hexadecanol |  |  |
| o-Xylene |  |  |

**Table 3.** Compounds were detected in Melia and its corresponding control samples. From here we see that endophytes are well-equipped to make their own compounds, and such compounds can be different from what is made by the host plant.

| **Melia + endophyte** | **Endophyte** | **Shared** |
| --- | --- | --- |
| (9Z)-Hexadecenoic acid | (2E)-Pentenal | 1-Docosene |
| (E)-Caryophyllene | (3E)-Octadecene | 1-Heptadecene |
| 1,19-Eicosadiene | 2,6,10,15-tetramethyl heptadecane | 1-Hexadecene |
| 1,2,3,4-Butanetetrol, [S-(R*,R*)]- | 2-Methyl-2-docosene | 1-Octadecene |
| 1,3-bis(1,1-dimethylethyl)-benzene | 2-Nonanol | 11-Methyldodecanol |
| 1-Decene | 2-methoxy-3-methyl phenol | 2,4-bis(1,1-dimethylethyl) phenol |
| 1-Dodecene | 2-methyl-, 3-methylbutyl propanoate | 2,5-bis(1,1-dimethylethyl)-phenol |
| 1-Eicosene | 2-methyl-heptadecane | 2,6,10,14-tetramethyl hexadecane |
| 1-Hexacosene | 3,5-dimethoxy phenol | 2,6,10,14-tetramethyl pentadecane |
| 1-Tetradecene | 3-hydroxypropyl octadecanoate | 2,6,10-trimethyl pentadecane |
| 1-methoxy-2-methylbutane | 5-Decanone | 2,6,11-trimethyl dodecane |
| 13Z-Docosenamide | 6-Dodecanone | 3-methyl-2-butenal |
| 2,3-Butanediol | 9,19-Cyclolanost-25-en-3-ol, 24-methyl-, (3.beta.,24S)- | 3-methylbutyl butanoate |
| 2,3-Butanediol, [R-(R*,R*)]- | Ethyl Oleate | 4-methyl-pentadecane |
| 2,3-Butanediol, [S-(R*,R*)]- | Geranyl isovalerate | 6,10,14-trimethyl-2-pentadecanone |
| 2,5-dimethyl-2,5-hexanediol | Hentriacontane | 6-Tridecanone |
| 2,5-dimethyltetradecane | Heptacosane | 7,9-Di-tert-butyl-1-oxaspiro(4,5)deca-6,9-diene-2,8-dione |
| 2,6,10-trimethyl tetradecane | Methyl -9Z-hexadecenoate | C20-ol |
| 2,6-dimethyl-2,6,10-undecatrien-8-ol | Methyl pentadecanoate | Docosane |
| 2,6-dimethyl-nonane | Octacosanol | Dodecane |
| 2-Acetylcyclopentanone | Octadec-9-enoic acid | Eicosane |
| 2-Pentanol |  | Eicosanoic acid |
| 2-hydroxy-1-(hydroxymethyl)ethyl esterhexadecanoic acid |  | Ethanone, 1-(9-anthracenyl)- |
| 2-hydroxy-3-methyl butanoic acid |  | Ethyl (9E)-octadecenoate |
| 2-hydroxyethylpropanoate |  | Ethyl 13-methyl-tetradecanoate |
| 2-methyl tridecane |  | Ethyl hexadecanoate |
| 2-methyl undecane |  | Ethyl octadecanoate |
| 2-methyl-1,3-butanediol |  | Ethyl propanoate |
| 2-methylbutanoic acid |  | Heneicosane |
| 2-methylpropyl ethanoate |  | Heptadecane |
| 3,3-dimethyl hexane |  | Hexacosane |
| 3,5-Dimethyldodecane |  | Hexadecane (C16) |
| 3,7-dimethyl-decane |  | Isopropyl myristate |
| 3-Penten-2-ol |  | Isopropyl tetradecanoate |
| 3-methyl tridecane |  | Methyl octadecanoate |
| 4',6'-Dihydroxy-2',3'-dimethylacetophenone |  | Nonadecane |
| 4,6-dimethyl dodecane, |  | Octacosane |
| 4-Methyl-5-nonanone |  | Octadecane (C18) |
| 4-Pinol |  | Octadecanoic acid |
| 4-methyl dodecane |  | Oleic Acid |
| 4-methyl hexadecane |  | Pentacosane |
| 4-methyl-2,6-Octadiene-4,5-diol |  | Pentadecane |
| 4-methyl-3-hexanol, |  | Pentadecanoic acid |
| 5,8-Tridecadione |  | Squalene |
| 5-Tetradecanol |  | Styrene |
| 5.alpha.-Spirostan-24-ol,<br>(24R,25S)- |  | Tetracosane |
| 6-Pentadecanone |  | Tetradecane |
| 8,11,14-Eicosatrienoic acid,<br>(Z,Z,Z)- |  | Triacotane |
| 8,11,14-Eicosatrienoic acid,<br>methyl ester |  | Tricosane |
| 8-methyl heptadecane |  | Tridecane |
| 9,12-Octadecadienoic acid<br>(Z,Z)- |  | Z,Z-10,12-Hexadecadienal |
| 9-Octadecenamide, (Z)- |  | n-Hexadecanoic acid |
| 9-Octadecenoic acid, (E)- |  | n-Hexadecanol |
| 9-Oxa-bicyclo[3.3.1]nonane-2,6-dione |  | n-Pentadecanol |
| 9-Tricosene, (Z)- |  | n-Tetracosanol-1 |
| A'-Neogammacer-22(29)-ene |  |  |
| Acetoin |  |  |
| Benzoic acid |  |  |
| Benzyl alcohol |  |  |
| Butanedioic acid |  |  |
| Butanoic acid |  |  |
| C14-ol Tetradecanol |  |  |
| Cis-3-methylpent-3-ene-5-ol |  |  |
| Decane, 2,3,5-trimethyl- |  |  |
| Dimethyl tetradecanedioate |  |  |
| Dodecane |  |  |
| Ergosta-4,6,8(14),22-tetraen-3-one |  |  |
| Ergosta-7,22-diene-3,5,6-<br>triol,<br>(3.beta.,5.alpha.,6.beta.,22E)- |  |  |
| Ergosterol |  |  |
| Ethyl (9Z)-octadecenoate |  |  |
| Ethyl 9-hexadecenoate |  |  |
| Ethyl heptadecanoate |  |  |
| Ethylbenzene |  |  |
| Glycerin |  |  |
| Heptadecanoic acid |  |  |
| Heptane |  |  |
| Hexanal |  |  |
| Hexanoic acid |  |  |
| L-Lactic acid |  |  |
| Limonene |  |  |
| Lup-20(29)-en-3-one |  |  |
| Methyl 9Z-Octadecenoate |  |  |
| Methyl dodecanoate |  |  |
| Methyl hexadecanoate |  |  |
| Methyl linoleate |  |  |
| Methyl tetradecanoate |  |  |
| Naphthalene, 1,2,3,4-tetrahydro-1-phenyl- |  |  |
| Neoergosterol |  |  |
| Nonanal |  |  |
| Octane |  |  |
| Pentadecanal |  |  |
| Phenol |  |  |
| Phenyl ethyl alcohol |  |  |
| Phytol |  |  |
| Solanidan-3-one |  |  |
| Tetracosane |  |  |
| Tetradecanoic acid |  |  |
| Toluene |  |  |
| Tridecane |  |  |
| Undecane |  |  |
| Undecane, 2,5-dimethyl- |  |  |
| Undecanoic acid |  |  |
| Z,E-2,13-Octadecadien-1-ol |  |  |
| alpha-trans-Bergamotene |  |  |
| cis-10-Heptadecenoic acid |  |  |
| n-Decanoic acid |  |  |
| o-Xylene |  |  |
| α-Copaene | | |
| α-Longipinene | | |

## Supporting information

raw PCA datasets

